# Longitudinal profiling of the pregnancy plasma proteome through organic solvent precipitation and nano LC-MS/MS

**DOI:** 10.1101/2023.12.11.571058

**Authors:** A.L George, E Cook, G.C.S Smith, D.S Charnock-Jones, S O’Rahilly, F Reimann, F.M Gribble, R.G Kay

## Abstract

Proteins secreted from maternal, fetal, and placental tissue are vital for processes like placental development, immunotolerance, and fetal growth, and are associated with pregnancy complications, necessitating predictive biomarkers. In this study, we introduce an acetonitrile-based precipitation coupled with solid-phase extraction that addresses limitations of current low throughput blood-based biomarker discovery workflows. Our method is efficient and cost-effective, and identified 433 protein groups, showing specific tissue associations and enrichment in reproductive tissues such as the placenta, breast, and endometrium. Significant alterations in proteins related to hormonal regulation, immune modulation, and placental development throughout gestation were observed. This approach offers comprehensive insights into the circulating pregnancy proteome, but also provides a scalable solution for larger studies for biomarker discovery in the context of pregnancy complications.

**TOC:** 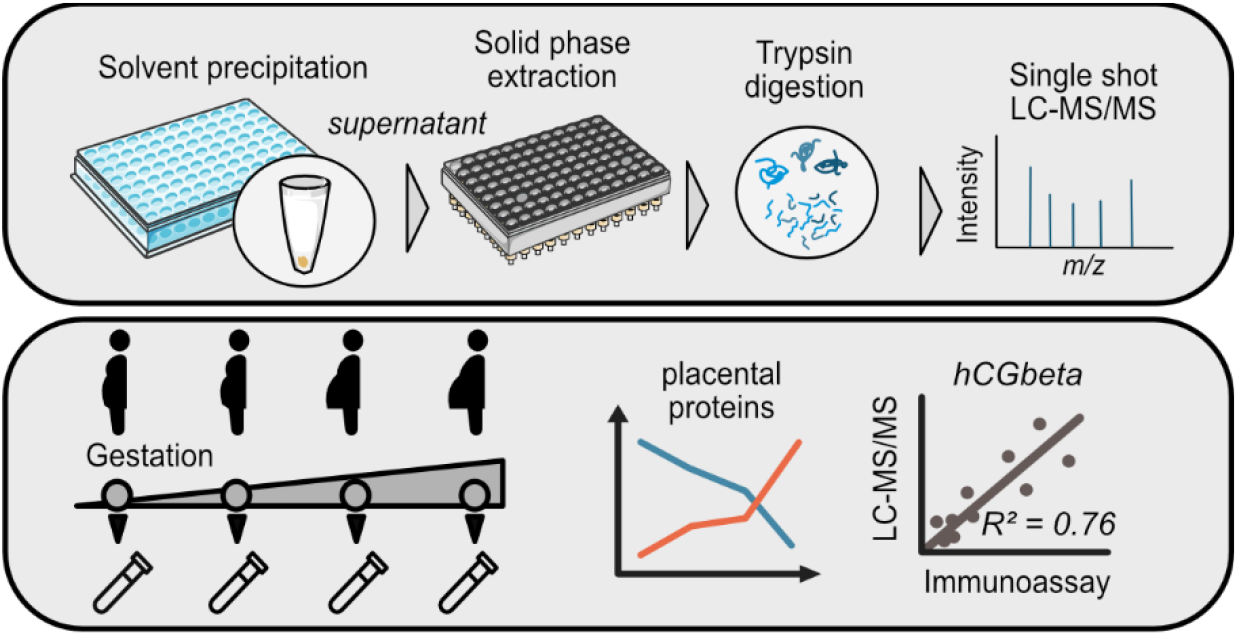

## Introduction

Proteins, including hormones, growth factors, cytokines, cell-adhesion molecules and immunomodulators play a crucial role in establishing and maintaining pregnancy. They promote fetal growth and development and orchestrate adaptations in maternal physiology to support successful gestation.^1, 2^ Their regulated secretion from fetal-placental and maternal tissue into the bloodstream supports placental development, implantation, immunotolerance and fetal growth.^3^ Importantly, perturbations in these physiological processes can lead to conditions such as preeclampsia, fetal growth restriction and preterm birth, which can arise without clear predictive biomarkers, necessitating extensive research for new biomarkers to enable a more personalised approach to pregnancy care. ^4-6^

Blood (plasma or serum) samples, collected during pregnancy provide a minimally-invasive and protein-rich source of biomarkers secreted from maternal, fetal and placental tissues.^7^ In recent years, these samples have been analysed with a focus on gaining a deeper understanding of the complex molecular adaptations key for healthy pregnancy, as well as to investigate indicators of placental function, fetal, and maternal health associated with pregnancy complications.^8^ Various blood-based biomarkers have been proposed to stratify the risk of different pregnancy complications but have struggled to provide the necessary sensitivity and specificity for early risk prediction.^9-11^

Advancements in proteomic technology for multiplexed analysis have accelerated capabilities in biomarker discovery. Explorative studies of protein abundance during both healthy and complicated pregnancies have adopted antibody or aptamer-based approaches, enabling sensitive detection of multiple proteins simultaneously.^12-14^ For example, Wang et al. explored a panel of 92 inflammatory markers using the proximity extension assay, O-Link platform, to investigate differences between patients with preeclampsia and healthy pregnant controls, as inflammation is reported to be involved in the occurrence and development of preeclampsia. They identified cysteine-cysteine motif chemokine ligand 20 (CCL20) as being significantly elevated.^15^ However, these platforms are limited by their capacity to measure only a predetermined set of proteins.

Liquid chromatography coupled with high-resolution mass spectrometry (LC-MS) offers distinct advantages for biomarker discovery and clinical proteomics compared to affinity-based approaches, enabling untargeted analysis with exceptional analytical specificity. Peptide sequences with a single-mass unit difference can be differentiated, allowing quantification of proteoforms and post-translational modifications.^16, 17^ Yet, analysis of plasma or serum samples is analytically challenging due to their high dynamic range of protein concentration, which is exacerbated by the presence of highly abundant proteins such as albumin, masking detection of the more relevant but less abundant putative biomarkers. Existing workflows for undepleted plasma analysis that can measure over 300 abundant circulating proteins in a 3-hour analysis, have not been adopted for identifying biomarkers of pregnancy complications.^18^ This is because signalling molecules likely to be markers of placental function circulate at low concentrations. Rather, studies of the circulating pregnancy proteome utilise in-gel digestion, extensive peptide fractionation or immunodepletion to remove the most abundant proteins and reduce sample complexity prior to mass spectrometry.^19, 20^ However, these approaches are costly, time consuming and introduce experimental variability.^21^ To overcome these challenges, a more efficient extraction approach is needed to eliminate sample preparation as a rate-limiting step and facilitate larger studies with appropriate sample sizes for verifying proposed markers.

Organic solvent precipitation is a simple, cost-effective and efficient approach for removing high abundance and high molecular weight proteins.^22^ This method, applicable to animal plasma as demonstrated in murine pregnancy studies^23, 24^, is capable of enriching for low molecular weight proteins circulating at much lower concentrations. In this study, we employed an acetonitrile-based precipitation method coupled with solid-phase extraction to rapidly prepare plasma for mass spectrometry analysis. By precipitating the most abundant circulating proteins, we measured a plethora of proteins secreted from the placenta and other reproductive tissues, that significantly change in abundance during uncomplicated pregnancy in healthy controls at 12-, 20-, 28- and 36-weeks gestational age. Samples were obtained from the Pregnancy Outcome Prediction (POP) study which was a prospective sample collection including greater than 4000 participants from the Cambridge area in the United Kingdom.^25^

## Materials and methods

### Study design and sample collection

The POP study recruited only nulliparous women, upon confirmation of pregnancy after a dating scan. Participants enrolled into the study had blood taken at approximately 12, 20, 28 and 36 weeks gestational age (wkGA). The POP study was ethically approved by the Cambridge Local Research Ethics Committee 2 and written informed consent obtained from all participants. Both plasma and serum were collected at each visit, with the serum samples used for protein quantitation on the Cobas e411 clinical analyser. The plasma was collected into EDTA tubes, centrifuged at +4ºC, separated into 4 aliquots and then stored at −80ºC until required. Plasma samples at four time points from six healthy volunteers with uncomplicated pregnancy were obtained for this prospective LC-MS/MS study.

### Sample preparation

Plasma samples (25 *μ*L) were diluted with 25 *μ*L of 6M guanidine hydrochloride and mixed on an Eppendorf MixMate for 1 minute at 1500 RPM. Proteins were precipitated with 300 *μ*L of 75% ACN in water with 50 ng/mL of bovine insulin as an internal standard. The plate was mixed for 1 minute at 1500 RPM, before centrifugation at 3900 x g for 10 minutes at +4ºC. The supernatant was transferred to an Eppendorf Lo-Bind 96 well plate and evaporated under oxygen free nitrogen (OFN) heated at 40 ºC. The dried sample residue was reconstituted into 200 *μ*L of 0.1% formic acid in water and transferred to a Waters HLB PRiME *μ*-elution SPE plate and pushed through carefully under positive pressure OFN. The wells were washed with 200 *μ*L of 0.1% formic acid in water, then 200 *μ*L of 5% methanol in 1% acetic acid in water. The proteins were eluted with 2 x 30 *μ*L of 60% methanol in 10% acetic acid in water into a fresh Lo-Bind plate and the eluant dried under OFN at 40 ºC. To reduce and alkylate disulphide bonds, 75 *μ*L of 10 mM DTT in 50 mM ammonium bicarbonate was added and the plate incubated at 60 ºC for 1 hour before addition of 20 *μ*L of 100 mM iodoacetamide in 50 mM ammonium bicarbonate and left in the dark for 30 minutes at room temperature. Samples were left on the bench at RT in the light for 20 minutes before 10 *μ*L of 100 *μ*g/mL trypsin gold in 50 mM ammonium bicarbonate was added and incubated overnight at 37 ºC. The digestion was halted by the addition of 20 *μ*L of 1% formic acid in water and 10 *μ*L was injected for LC-MS/MS analysis.

### Liquid chromatography mass spectrometry (LC-MS/MS)

Samples were analysed using a Thermo Fisher Ultimate 3000 nano LC system coupled to a Q Exactive Plus Orbitrap mass spectrometer (ThermoScientific, San Jose, CA, USA). Tryptic peptides were loaded onto a 0.3 x 5mm peptide trap column (ThermoFisher Scientific) at 30 *μ*L per minute for 15 minutes before switching in line with a 0.075 × 250 mm nano easy column (ThermoFisher Scientific) flowing at 300 nL/min. Initial gradient conditions were 2% ACN rising to 50% over 90 minutes, returning to initial conditions after a 20-minute wash at 90% ACN, giving a total run time of 130 minutes. Electrospray analysis was performed using a spray voltage of +1.8 kV, and a full scan range of 400–1600 m/z was performed at a resolution of 75,000 before the top 10 ions of each spectrum were selected for MS/MS analysis. Existing ions selected for fragmentation were added to an exclusion list for 30 s.

### Data analysis

Acquired mass spectrometry data was searched against the human Uniprot protein database (downloaded Aug 2021) using PEAKS 8.5, with fixed carbamidomethylation on cysteine residues, and variable modifications for oxidised methionines and asparagine / glutamine deamidation. Results were filtered to include only proteins with a 1% false discovery rate (FDR) and at least 1 unique peptide (Table S1A). Tissue enrichment analysis was performed with TissueEnrich (https://tissueenrich.gdcb.iastate.edu/). Peptides from proteins related to pregnancy (selected based on their number of spectral matches, and sensitivity and selectivity for that protein) were targeted for label free quantitation (manual integration) and their peak areas were calculated using the data raw files in Quan Browser (ThermoScientific, San Jose, CA, USA)(Table S1B). For global analysis, peak areas were Log2 transformed and filtered to include only proteins present in at least 4 of the 6 patient samples, in at least one time point. Missing values of this filtered dataset were then imputed in Perseus (version 2.0.1.1) to mimic proteins of low abundance that were below the limit of detection during some timepoints (width 0.3, downshift 0.1). All data were expressed as a ratio to gestational age week 12. Gene Ontology enrichment analysis of proteins significantly changing during gestation was performed using the bioinformatic resource DAVID.^26^

### Automated immunoassay analysis

During the POPs study, maternal serum was analysed for alpha fetoprotein (AFP), chorionic gonadotropin (hCG), Pregnancy associated plasma protein-A (PAPP-A), placental growth factor (PlGF), and soluble FMS like Tyrosine kinase 1 (sFlt-1) on a Cobas e411 analyser (Roche Diagnostics, Mannheim, Germany).^27^ The serum hCG concentrations of the samples from the six individuals used in this study were compared against the respective LC-MS/MS derived values.

### Statistical analysis

Statistical analysis was performed in R (v4.0.4). One-way ANOVA repeated measures was used to investigate differences in protein abundance associated with gestational age, followed by pairwise t-tests for multiple comparison of gestation weeks (FDR adjusted p-value < 0.05).

## Results and discussion

### Characterisation of the circulating proteome

To profile the longitudinal plasma proteome during healthy pregnancy, maternal plasma samples were collected at approximately 12, 20, 28 and 36 wkGA (Figure 1A). A central challenge with plasma or serum-based proteomics is the limitation of achievable sensitivity, as the prevalence of highly abundant proteins such as albumin dominate the measure of quantified peptides by mass spectrometry, making the detection of less abundant biomarkers originating from disease tissue much more difficult.^29^ Removal of these highly prevalent proteins is therefore necessary, but immunodepleting or fractionation approaches require expensive reagents and lengthy workflows, which are less than optimal for a high-throughput workflow to accommodate the large sample sizes that are necessary for clinical biomarker studies. Protein precipitation using organic solvents offers an alternative method for protein enrichment, and can be rapidly performed in 96-well plate format, is cost-effective as there is no need for specialised reagents or antibodies, and reliably detects low-concentration plasma proteins.^22, 23^ In this study, we diluted plasma with guanidine hydrochloride to dissociate small proteins bound to blood carrier proteins, like insulin-like growth factor 1 (IGF-1) which circulates bound to IGF binding protein 3. Subsequently acetonitrile (75% in water) was used to precipitate high abundance proteins but retain the soluble, less abundant proteins in the supernatant. The supernatant was evaporated and subjected to solid-phase extraction to remove unwanted hydrophobic small molecules, followed by in-solution digestion, and was analysed by label-free single-shot mass spectrometry (Figure 1B).

**Figure 1.**
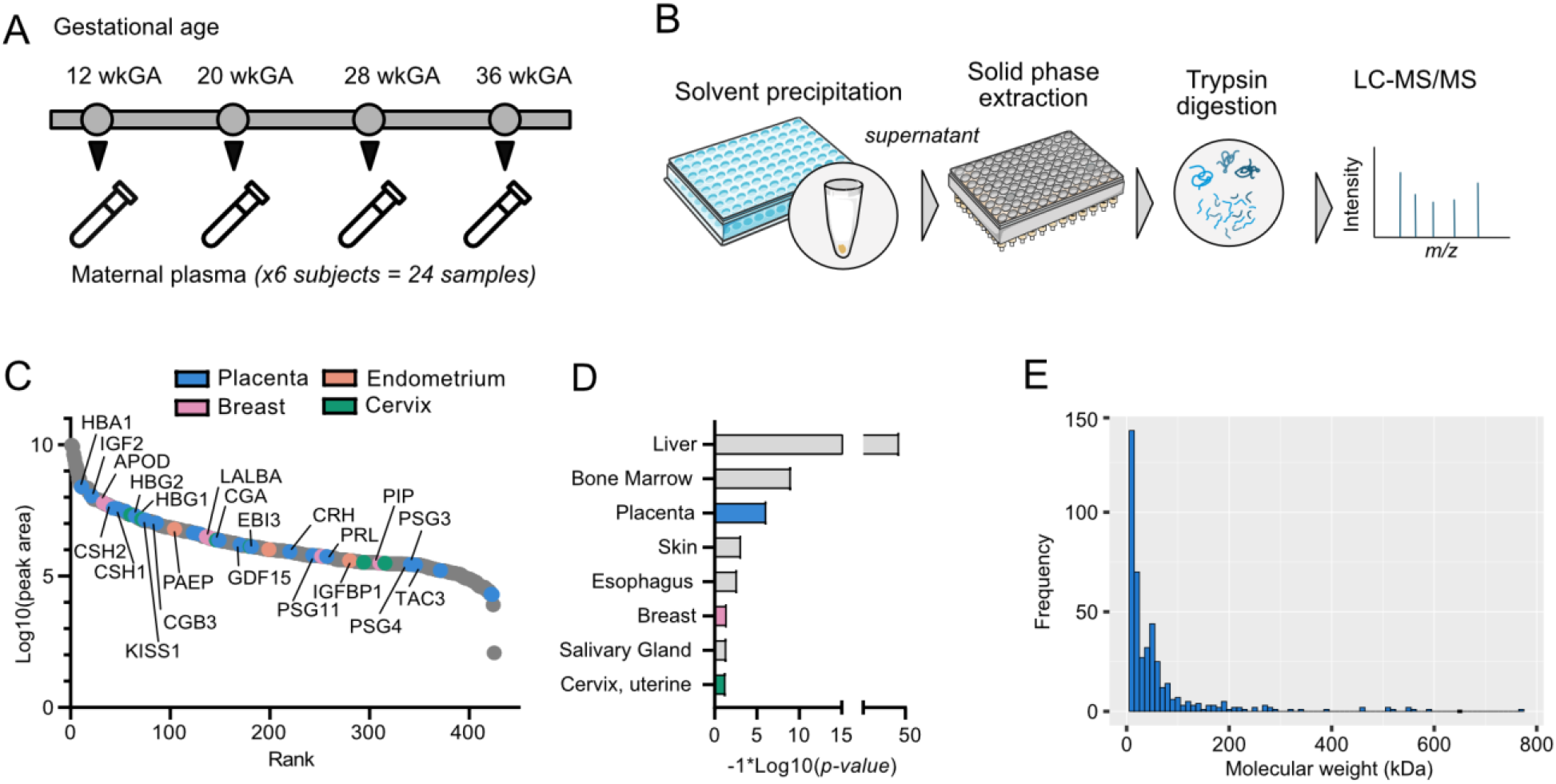
Characterisation of the circulating proteome during pregnancy. **A)** Sample collection time points of the six subjects. **B)** Sample preparation workflow to isolate the circulating proteome. Plasma proteins are diluted with guanidine hydrochloride before precipitation with acetonitrile and solid phase extraction. Extracted proteins are digested with trypsin and peptides analysed by LC-MS/MS. **C)** Ranked average abundance plot of all proteins in maternal plasma demonstrating depletion of abundant circulating proteins, enriching for less abundant signalling proteins. **D)** Tissue enrichment analysis of all proteins detected revealed a significant enrichment in placental (blue), breast (pink) and uterine (green) proteins. **E)** Histogram showing frequency in molecular weight of the identified 433 plasma proteins.

A total of 433 protein groups were identified, exhibiting specific tissue associations, as indicated in Figure 1C and Figure 1D. The acetonitrile precipitation method effectively depleted albumin, although other abundant plasma proteins such as low molecular weight apolipoproteins (APOA2, APOC3, APOA1, APOC2) that are secreted from the liver and circulate at *μ*g/mL to mg/mL levels were also measured and ranked highest when ordering each identified protein by their average intensities across all samples (Figure 1C). Tissue enrichment analysis highlighted many proteins that are significantly enriched in the liver, an expected result, given the liver’s major role in the biosynthesis of many plasma proteins. Notably, many of the other identified proteins are significantly enriched in female reproductive tissues, including the placenta, breast, cervix, or endometrium (Figure 1D).

For example, 24 proteins with elevated expression in the placenta were identified, thirteen of which are classified as tissue enriched, defined by the Human Protein Atlas to have a four-fold higher expression in the placenta compared to other tissues (Figure S1A). They included pregnancy-specific β-1-glycoproteins (PSG11, PSG3 and PSG4) that are uniquely produced by placental syncytiotrophoblast cells and play key roles in immunomodulation.^30^ A recent mass spectrometry-based plasma proteomics study proposed 16 candidate biomarkers for ectopic pregnancy prediction, which included PSG11 and PSG3, but their workflow consisted of a two-step depletion of abundant plasma proteins with IGY-14 and super mix immune-depletion columns, followed by in-gel fractionation (6 fractions) to achieve such proteomic depth.^31^

Compared to other methods, our single-shot analysis, with minimal sample handling, offers sufficient sensitivity to detect low abundance proteins known to circulate in maternal blood at low ng/mL range such as leptin and tachykinin-3. Leptin, for instance, is produced by both maternal and fetal adipose tissue, but also by the placenta to support implantation, human chorionic gonadotrophin production, placental growth, amino acid uptake, and mitogenesis.^32^ Perturbations in leptin levels have been correlated with various reproduction and gestation disorders such as polycystic ovary syndrome, recurrent miscarriage, gestational diabetes mellitus, pre-eclampsia, and intrauterine growth restriction.^33^ The neuropeptide, tachykinin-3 (TAC3), has enriched expression in the outer syncytiotrophoblast of the placenta and was among the lowest ranking proteins in the abundance plot of circulating proteins (Figure 1C). During normal pregnancy, it circulates at 0.29–0.61 nmol/L and is elevated in women suffering with preeclampsia.^33, 34^ Identifying these low abundance proteins alongside abundant apolipoproteins and immunoglobulins, demonstrates the method’s ability to measure proteins over a wide dynamic range.

Plotting the frequency of protein molecular weight demonstrates that precipitation using 75% acetonitrile effectively removes large molecular weight proteins and enriches those with a molecular weight of less than 60kDa (Figure 1E). In another study, a different ratio of blood to the organic solvent (1:2 ratio, 100% acetonitrile, v/v) was employed which identified mainly 2.5 kDa proteins, along with some in the range of 2.5 to 10 kDa, emphasising the adaptability of the extraction approach depending on the proteins of interest.^35^

Peak areas of peptides from proteins with known roles in pregnancy were manually integrated and plotted by gestational age (Figure 2). This highlighted differences in the low molecular weight proteome and bioactive peptide profiles between individuals. For example, cystatin (13kDa), trefoil factor (7kDa), insulin-like growth factor 1 (IGF-1) (8kDa) and insulin-like growth factor 2 (IGF-2) are all low molecular weight proteins that regulate or support processes such as immune modulation, implantation, or fetal growth. Plasma levels of cystatin-C during normal pregnancy are described as stable during the first trimesters of pregnancy, but increase significantly during the final trimester.^36^ These patterns were also identified in our dataset, as cystatin-C increased over pregnancy in all participants (Figure 2A). Cystatin-C plasma levels have been explored as a marker of renal function in preeclampsia.^37^ Trefoil factors are peptides secreted from mucosal surfaces for protection and repair, primarily in the gastrointestinal tract. During pregnancy, circulating levels of TFF3 are reported to be 47 times higher than postpartum^38^, and were significantly elevated in our dataset (Figure 2B). Growth factors, insulin-like growth factor 1 (IGF-1) and insulin-like growth factor 2 (IGF-2) that play important roles in cell growth, differentiation, and development, are involved in the development of the placenta and fetal tissues during pregnancy^39^ and were significantly higher compared to week 12 (Figure 2 C and D).

**Figure 2.**
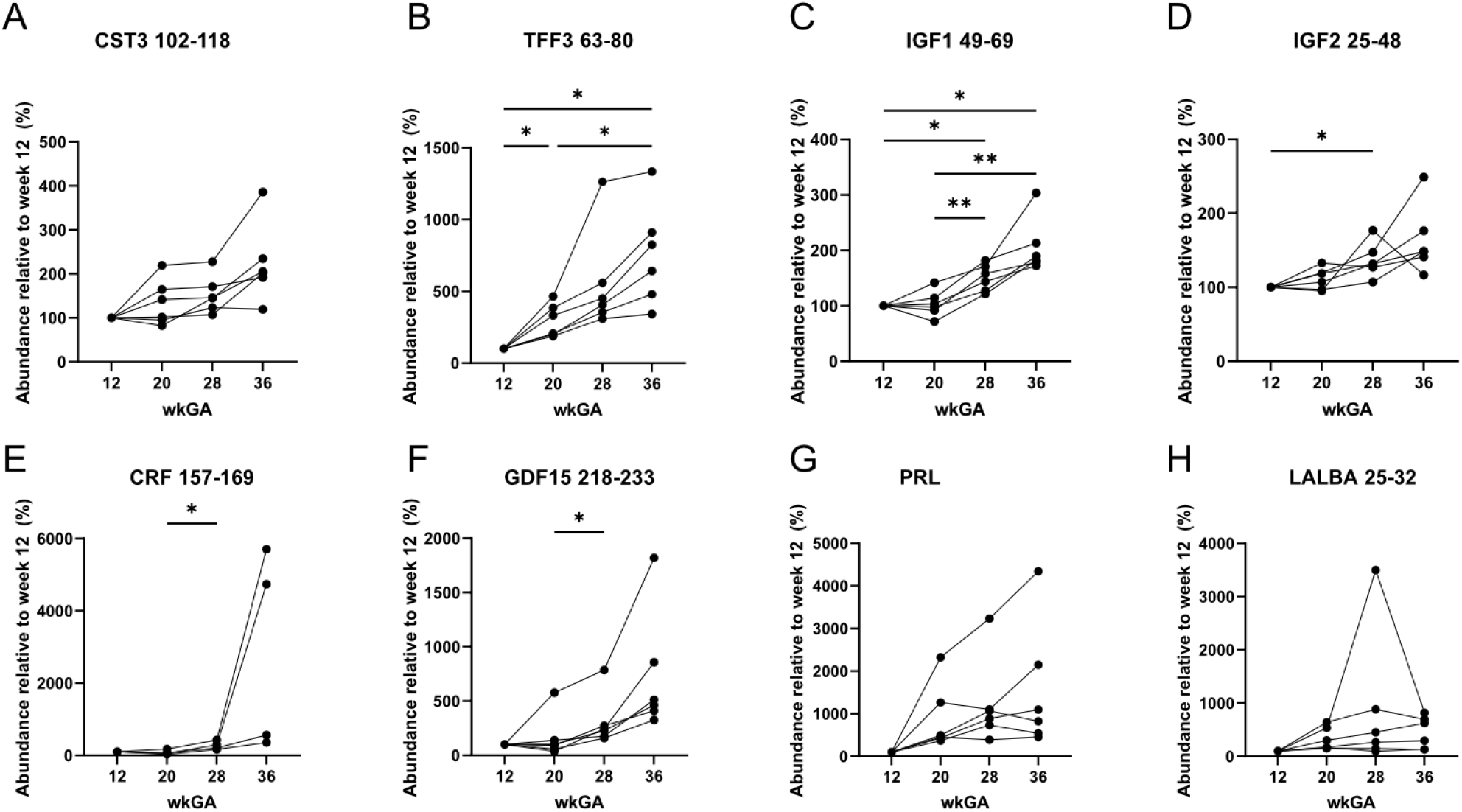
Peak area of peptides from pregnancy-associated proteins relative to week 12 gestational age. Numbers correspond to the amino acid positions of the monitored peptide in the primary amino acid sequence. **A)** cystatin-C (CST3), **B)** trefoil factor 3 (TFF3), **C)** insulin-like growth factor I (IGF-I),**D)** insulin-like growth factor II (IGF-II), **E)** corticoliberin (CRF), **F)** growth-differentiation factor 15 (GDF15), **G)** prolactin (PRL), **H)** alpha-lactalbumin (LALBA).

Corticoliberin, also known as corticotropin releasing hormone (CRF) is a peptide hormone secreted into the blood at very low levels by the hypothalamus but is also highly expressed in the placenta. Placental CRF secretion increases exponentially, peaking at birth and maternal plasma CRF levels increase accordingly from the second trimester. CRF levels in maternal plasma in the first and second trimester have been measured to remain within non-pregnant range (20pg/mL at 28 weeks) but increase significantly in the final trimester to 1320 pg/mL at 40 weeks triggering parturition^40, 41^, and high maternal levels have been associated with preterm labour.^42^ In our dataset the peptide was detected in four of the six volunteers (Figure 2E). CRF has been identified previously by LC-MS approaches in placental tissue using solid phase extraction, in-gel digestion or a molecular-weight cutoff filter^43^, but is reported only once before to be detected in plasma using an LC-MS method that included immunodepletion and extensive fractionation.^20^

GDF15, a member of the TGF-beta family, exhibits high expression in the placenta, and in healthy pregnancy circulating levels rise rapidly in maternal blood during the first trimester and continue to elevate until term, as shown with our dataset (Figure 2F). Importantly, immunoassay measurement of GDF15 in human plasma and serum has been reported to underestimate levels in individuals who carry a histidine (H) to aspartate (D) variant at position 202 in the propeptide.^44^ However, mass spectrometry can distinguish between these variants and can therefore be utilised to gain an understanding of the origin of proteins that circulate in maternal blood. By exploiting the specificity of mass spectrometry, the stoichiometry of the H and D peptides was measured to determine the relative levels of maternal or fetal derived GDF15 in the maternal circulation, which confirmed circulating GDF15 in human pregnancy is predominantly of fetal origin.^45^

Prolactin (PRL) is a growth hormone which acts primarily on the mammary gland to stimulate lactation. Additionally, it plays important roles in metabolism and immune system regulation.^46^ In healthy non-pregnant women, circulating prolactin levels are low but rise during gestation (Figure 2G). Moreover, alpha-lactalbumin (LALBA), a human milk protein, that is expressed in the glandular and ductal cells the breast. As a result of breast development during pregnancy, LALBA levels rise significantly and due to tissue leakage, its presence can be detected in the blood at increasing levels. Studies have indicated a significant increase until the mid-trimester, after which concentrations remained stable until term^47^, a trend reflected in our data for five of the participants (Figure 2H).

### Changes in the circulating proteome associated with gestational age

For unbiased assessment of the method for measuring significant changes in protein abundance during pregnancy, we compared protein group abundance at each of the four time points by one-way repeated measures ANOVA. As the expression of proteins can change considerably over gestational weeks and circulating concentrations can vary hundred-fold, missing values were imputed to mimic proteins of low abundance that were below the limit of detection during some timepoints, resulting in comparison of 231 protein groups. This identified 16 proteins with significant changes associated with gestational age (Figure 3A)(Table S1C and Table S1D). These protein groups exhibited distinct profiles as either increasing (red) or decreasing (blue) during pregnancy. Human chorionic gonadotropin hormone (hCG) is a glycoprotein consisting of two non-covalently linked subunits of alpha and beta (hormone specific subunit). While the hormone is produced by trophoblast cells at high levels very early on during pregnancy (and indeed is used as the measured target of point of care pregnancy tests), it peaks at week 10 before gradually declining and stabilising until term. Isoforms of the hormone with identical amino acid sequences exist (hCG, sulfated hCG, hyperglycosylated hCG, hCG free ß-subunit and hyperglycosylated hCG free ß-subunit) and are produced at different concentrations, by different cells and with different functions. Different immunoassays vary in their ability to recognise these isoforms forms, although there is no evidence necessitating accuracy of its measurement to diagnose pregnancy or non-pregnancy.^48^ In our study, we detected beta subunits, CGB5, CGB7 and CGB2, as being highest in maternal plasma at week 12 of gestation, decreasing by week 20, then remaining constant during weeks 28 and 36 (Figure 3A). These measurements align with known secretion patterns during healthy pregnancy. Relative quantification for hCG over the course of pregnancy measured by LC-MS/MS showed good correlation (R2 = 0.76) with measurements from the Roche Cobas e411, which is a CE marked clinical analyser used in many hospital laboratories (Figure 3B).

**Figure 3.**
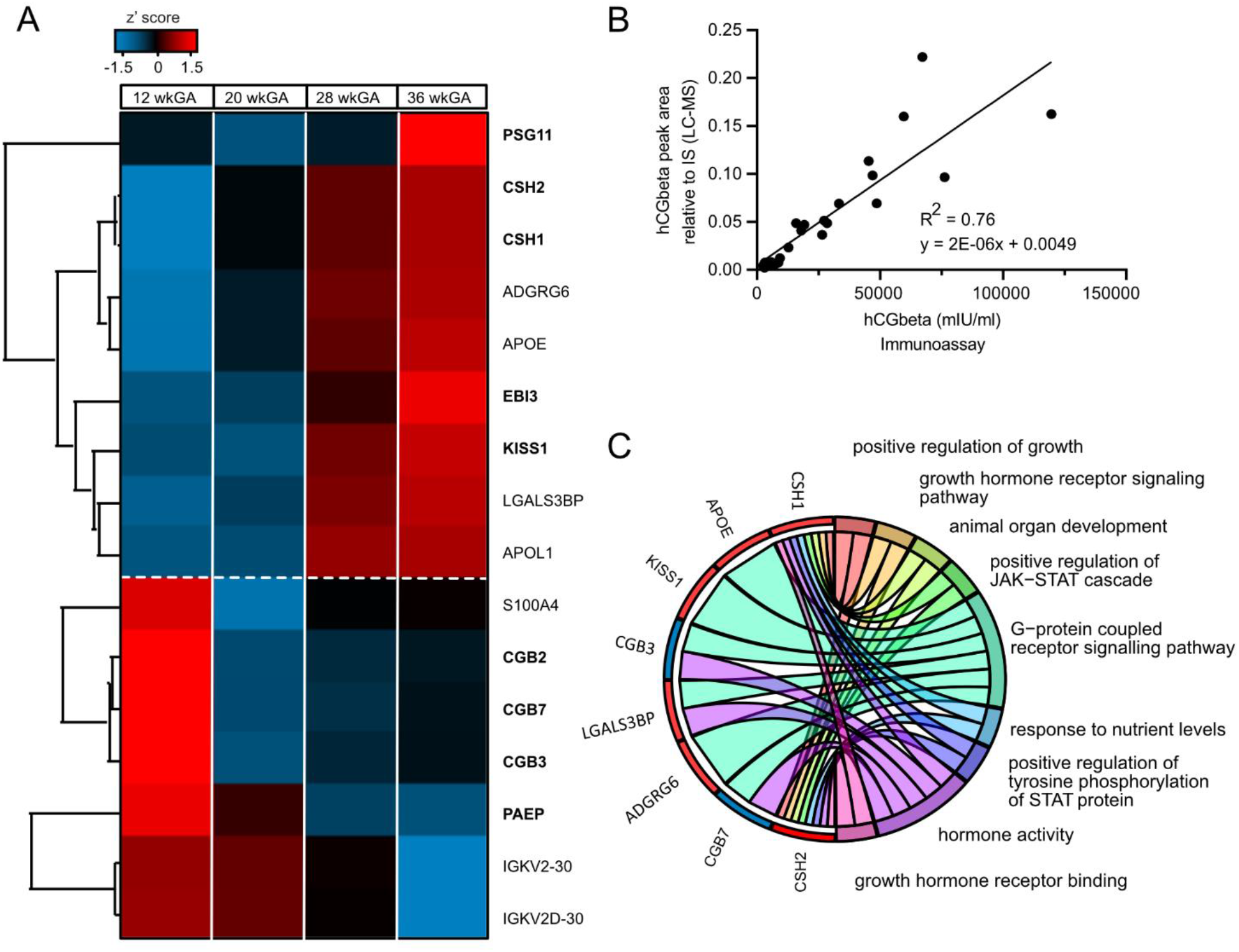
Changes in the circulating proteome associated with gestational age. **A)** Hierarchical clustering of proteins with significant changes in abundance associated with gestational age (ANOVA p < 0.05). Increased abundance is represented in the heat map in red and those decreased are represented in blue. Gestational age is shown. **B)** Correlation between hCGbeta levels measured relatively by LC-MS/MS compared with immunoassay quantification in all 24 samples. **C)** Chord diagram presenting gene ontology enriched terms in proteins that significantly change with gestational age.

Functional annotation through gene ontology (GO) enrichment analysis of the significantly changing proteins highlighted the importance of hormones in regulating gestation, as the molecular function, hormonal activity, was most significantly enriched (6.17E-07) (Table S1E). In addition to CGB5, CGB7 and CGB2, the growth factor placental lactogen significantly changed (CSH1 and CSH2). Placental lactogen, also known as chorionic somatotropin, is a peptide hormone produced by syncytiotrophoblast cells during human pregnancy. This growth hormone primarily circulates in the maternal blood, and is detectable from six weeks of gestation.^49, 50^ As seen in Figure 3C, levels increase throughout gestation, with levels thought to reflect total placental mass.^49^

Additionally, we noted changes in the endometrial protein, glycodelin (GD) also known as progestogen-associated endometrial protein (PAEP), which progressively declined in maternal plasma over the course of gestation (Figure 3A). Studies have shown concentrations in maternal serum to increase in the case of premature rupture of membranes.^51^ This low molecular weight protein has been detected previously in maternal serum using LC-MS and was identified in the aforementioned Beer at al. study of ectopic pregnancy, but a lengthy depletion and enrichment protocol was previously adopted, and verification was limited because of the moderate sample size.^31^

We also observed significant changes in the interleukin-27 subunit beta (EBI3) (48kDa) between weeks 12 and 36 of gestation, reflecting the cytokine’s role in regulating maternal immune tolerance and angiogenesis during pregnancy.^52^ This protein has also been identified as a potential biomarker for preeclampsia, where the circulating plasma levels were higher in patients with preeclampsia (∼1 ng/mL) compared to healthy women (∼800 pg/mL).^53^

Kisspeptin is a peptide of approximately 6 kDa that stimulates release of hypothalamic gonadotropin-releasing hormone and has a role in regulating trophoblast invasion. It is expressed in various tissues but is primarily secreted by the early syncytiotrophoblasts, increasing levels in the maternal circulation thousand-fold during healthy pregnancy. Kisspeptin has been proposed as a biomarker for placental function, as it increases progressively over pregnancy. Our data mirrored this secretion pattern, as levels increased with gestational age. Low circulating levels are associated with placental dysfunction, which is linked to complications such as miscarriage, hypertensive diseases of pregnancy, fetal growth restriction, and gestational diabetes (GDM).^54^

In addition to placental-derived proteins, we observed changes in low molecular weight proteins with critical roles in supporting metabolic and nutritional adaptations during pregnancy. Proteins responsible for lipid transport and maintaining triglyceride homeostasis (APOL1, APOE) gradually increased with gestational age and were highest at week 36. Studies have suggested that levels of APOA2, APOC1, APOC3, and APOE are lower in women with gestational diabetes mellitus.^55^

While our plasma preparation method effectively addresses the challenge of highly abundant proteins that can dominate proteomic analyses, it predominantly enriches the low molecular weight proteome. Nonetheless, the proteins identified hold significant clinical relevance, playing crucial roles in biological processes vital for pregnancy, including implantation, placental development, and immune modulation, which are known to be perturbed during pregnancy complications. We also demonstrate the accuracy and quantitative performance of our method, as findings correlate well with immunoassay data and expected secretion patterns of known pregnancy-related proteins during normal pregnancy. In future applications, investigating these protein profiles during pregnancy complications holds promising clinical value, as many proteins identified have previously been proposed as biomarkers of pregnancy complication. Coupling this rapid extraction workflow with recent advances in mass spectrometry, such as faster scanning instruments capable of performing data independent acquisition experiments, supporting shorter analytical columns and gradients will significantly reduce run-times without compromising quantitative performance. This updated technology could accelerate biomarker discovery and verification in the context of pregnancy complications by enabling the analysis of larger sample cohorts for pregnancy-related clinical studies.

## Authorship contributions

ALG extracted, analysed the samples and performed the bioinformatics analyses. ALG and RGK prepared the manuscript, and all authors reviewed the manuscript. EC, GCSS, and DSCJ designed and led the POP study and supplied the samples. SOR, FMG, FR and RGK guarantee the study.

## Acknowledgments

The Prediction of Pregnancy Outcome Prediction Study (POPS) was supported by the Medical Research Council (United Kingdom; G1100221) and the National Institute for Health Research (NIHR) Cambridge Biomedical Research Centre (Women’s Health theme). The instrumentation used for the analysis was funded by the MRC Enhancing UK Clinical Research grant (MR/M009041/1). Research in the FMG/FR group is funded by MRC (MC_UU_00014/3) and Wellcome (220271/Z/20/Z), and the LC-MS peptidomics facility receives core support from MRC [MC_UU_00014/5] and Wellcome [100574/Z/12/Z]).

## Abbreviations

ANOVA: Analysis of variance
DAVID: The Database for Annotation, Visualization and Integrated Discovery
DTT: Dithiothreitol
GDM: Gestational Diabetes Mellitus
GO: Gene Ontology
OFN: Oxygen Free Nitrogen
RT: Room Temperature
wkGA: Week of Gestational Age

## Supporting information

***Figure S1***. *TissueEnrich analysis of detected proteins in maternal plasma*. ***Table S1A***. *Raw data of protein peak areas exported from PEAKS search*. ***Table S1B***. *Manual integration of peptides from protein with known pregnancy-related functions*. ***Table S1C***. *Proteins significantly associated with gestational age*. ***Table S1D***. *Pairwise comparison between weeks of gestational age*. ***Table S1E***. *Gene ontology enrichment analysis of significantly changing proteins*. ***Table S1F***. *LC-MS/MS file information, showing subject number and sampling time*.

