## Supporting information for "Longitudinal profiling of the pregnancy plasma proteome through organic solvent precipitation and nano LC-MS/MS"

Supplementary Figures

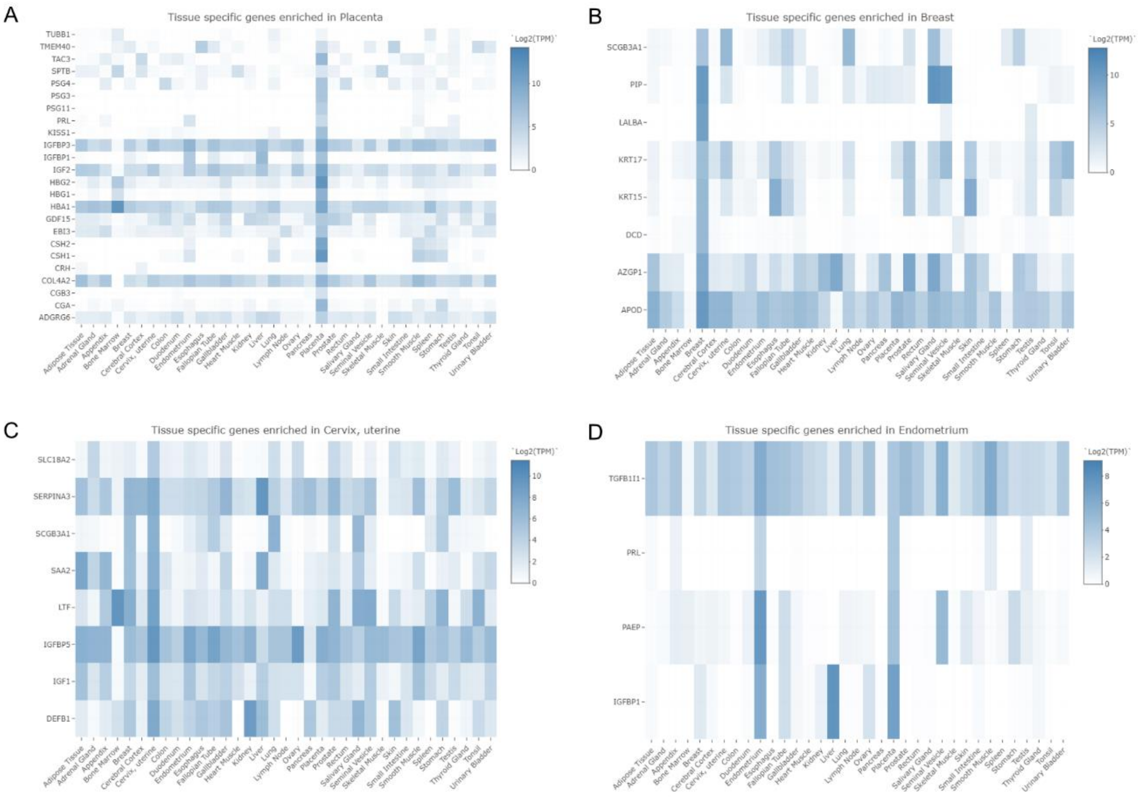

**Figure S1**. Tissue enrichment analysis. Proteins identified in maternal serum with elevated expression in the A) placenta, B) breast, C) cervix and D) endometrium.

**Supplementary Tables**

**Table S1A**. Raw data of protein peak areas exported from PEAKS search.

**Table S1B**. Data of manually integrated peak areas from peptides of select pregnancy-related proteins.

**Table S1C**. Proteins significantly associated with gestational age (one-way repeated measures anova)

**Table S1D**. Pairwise comparison between weeks of gestation.

**Table S1E.** Gene ontology enrichment analysis of proteins significantly associated with gestational age.

**Table S1F.** LC-MS/MS file information, showing subject number and sampling time.
